# Combining Mechanistic Models and Machine Learning for Personalized Chemotherapy and Surgery Sequencing in Breast Cancer

**DOI:** 10.1101/2020.06.08.140756

**Authors:** Cristian Axenie, Daria Kurz

## Abstract

Mathematical and computational oncology has increased the pace of cancer research towards the advancement of personalized therapy. Serving the pressing need to exploit the large amounts of currently underutilized data, such approaches bring a significant clinical advantage in tailoring the therapy. CHIMERA is a novel system that combines mechanistic modelling and machine learning for personalized chemotherapy and surgery sequencing in breast cancer. It optimizes decision-making in personalized breast cancer therapy by connecting tumor growth behaviour and chemotherapy effects through predictive modelling and learning. We demonstrate the capabilities of CHIMERA in learning simultaneously the tumor growth patterns, across several types of breast cancer, and the pharmacokinetics of a typical breast cancer chemotoxic drug. The learnt functions are subsequently used to predict how to sequence the intervention. We demonstrate the versatility of CHIMERA in learning from tumor growth and pharmacokinetics data to provide robust predictions under two, typically used, chemotherapy protocol hypotheses.

## I. Background

With over 70000 new cases in 2018 and almost 500000 prevalent cases on a 5-year prediction in Germany [1], breast cancer is still in the foreground of chronic diseases that require innovative therapeutic solutions. As the last decades have shown, early diagnosis and new drugs have led to impressive increases in survival rates of cancer patients. Yet, tailoring standard treatment schemes to patient needs is still a sought for objective. A personalised approach, requires new methods that exploit tumor biology and the effect chemotoxic drugs have upon the tumor, in order to sequence the interventions [2]. Neoadjuvant therapy (i.e., chemotherapy administered before surgery) has grown into a well-established, safe and often beneficial approach to breast cancer treatment. In terms of survival and overall disease progression, neoadjuvant and adjuvant treatments tend to be similar treatment choices for breast cancer [3]. Yet, the neoadjuvant treatment increases breast-conserving surgery levels and improves resectability by reducing the primary tumour. Moreover, it can support the early evaluation of the efficacy of the therapy chosen [4]. This assessment may allow the clinician to discontinue ineffective treatment or may help switch to another regimen to maximise response [5]. For a wide range of reasons, initial surgery accompanied by adjuvant chemotherapy may be the prevalent procedure or preferred choice for a particular patient compared with preoperative or neoadjuvant chemotherapy [6]. Moreover, patients who do not achieve a pathologic complete response after neoadjuvant chemotherapy, consider the use of adjuvant scheme [7]. Overall, it is reported that chemotherapy use in the neoadjuvant and adjuvant settings generally provides the same long-term outcome. But what is the best course of action for a particular patient? This question targets those quantifiable patient-specific factors (e.g. tumor growth curve and chemotherapy effect parameters, such as drug pharmacokinetics) that influence the sequencing of chemotherapy and surgery and taps into personalized therapy.

### A. Formalizing therapy sequencing

A model for personalized sequencing should include tumor cells growth and the effects of chemotherapy and surgery under cell-kill hypotheses. This hypothesis proposes that actions of chemotoxic drugs follow first order kinetics: a given dose kills a constant proportion of a tumor cell population (rather than a constant number of cells) [8]. Assuming that the the tumor size at time *t*_0_ = 0 is *V*_0_, there are two possible sequences:

- **Adjuvant chemotherapy** (chemotherapy after surgery). At time *t*_0_ *>* 0 a fraction of the tumor is removed through surgery and subsequently chemotherapy is administered with a killing rate of 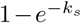 where *k*_*s*_ is a rate constant. The final size after the intervention, at *t*_*f*_ *> t*_0_ is *V*_*SC*_.
- **Neoadjuvant chemotherapy** (chemotherapy before surgery). At time *t*_0_ *>* 0 chemotherapy is administered with a predefined killing rate. At *t*_*f*_ *> t*_0_ a fraction 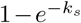 of the tumor is removed through surgery for a final size after the intervention *V*_*CS*_.

The question of interest in our study is if *V*_*SC*_ *> V*_*CS*_? If we consider *f* (*V*) the tumor growth model and *P* (*t, V*) the pharmacokinetics of the chemotherapeutic drug, we can formalize the two sequences as following:

- Sequence 1: Chemotherapy after surgery-before surgery

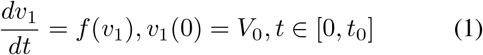

-after surgery

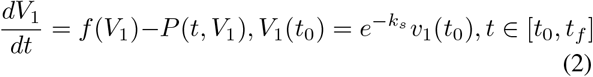

In this case, the final volume of the tumor is

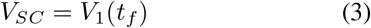
- Sequence 2: Chemotherapy before surgery-before chemotherapy onset

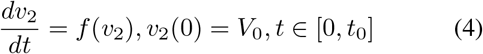

-after chemotherapy onset

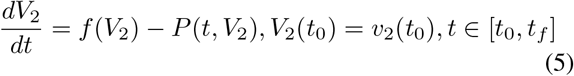

Hence, for the neo-adjuvant sequence, the final volume of the tumor is

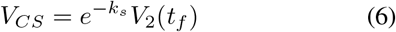

Such sequencing is typically valid under some cell-kill hypotheses. Basically, these hypotheses define the effect that the chemotherapy drug has upon the tumor. The two cell-kill hypotheses we employ in our study are the log-kill hypothesis [9] and the Norton-Simon hypothesis, respectively [10]. The log-kill hypothesis states that the effect the chemotoxic drug has upon the tumor is

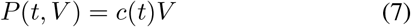

where *V* is the volume of the tumor or number of cells, and *c*(*t*) is a function proportional with the chemotoxic drug concentration at time *t*. On the other side, the Norton-Simon hypothesis, defines the effect of the chemotoxic drug as

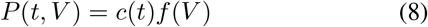

where *f* (*V*) is the tumor growth curve function depending on the volume of the tumor or number of cells. Various studies [11] considering average values over the populations of patients demonstrated that under the log-kill hypothesis *V*_*SC*_ *> V*_*CS*_ whereas under the Norton-Simpson hypothesis *V*_*SC*_ *> V*_*CS*_ or *V*_*SC*_ *< V*_*CS*_. In order to ensure that such sequencing is personalized, we explore how can a machine learning algorithm extract the two functions of interest, namely tumor growth function *f* and pharmacokinetics effect *P* from data, without constraining the choice of a specific model. Such a limiting approach would be detrimental for patients as it might not capture the tumor dynamics and the effect chemotherapy has for the long-term intervention.

### B. Models of tumor growth

A large variety of breast cancer tumor growth patterns were identified experimentally and clinically, and modelled over the years. Ordinary differential equations (ODE) tumor growth models [13] are typically used in cancer treatments planning. In our study, we explored three of the most representative and typically used scalar growth models, namely Logistic, von Bertalanffy, and Gompertz, described in Table I-B.

**TABLE I.**
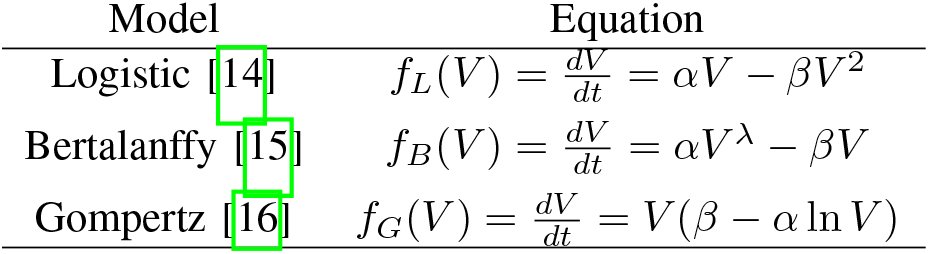
Overview of tumor growth models *f* (*V*) in our study. Parameters: *V* - volume (or cell population size through conversion - 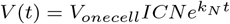 where *N* is the population size, *ICN* is the initial cell number, *V*_*onecell*_ is the volume of one cell and *k*_*n*_ is the rate constant for changes in cell number as considered in [17]), *α* - growth rate, *β* - cell death rate, *λ* - nutrient limited proliferation rate, *k* - carrying capacity of cells.

| Model | Equation |
| --- | --- |
| Logistic [14] | $f_L(V) = \frac{dV}{dt} = \alpha V - \beta V^2$ |
| Bertalanffy [15] | $f_B(V) = \frac{dV}{dt} = \alpha V^\lambda - \beta V$ |
| Gompertz [16] | $f_G(V) = \frac{dV}{dt} = V(\beta - \alpha \ln V)$ |

Despite their ubiquitous use, the aforementioned scalar tumor growth models are confined due to: a) the requirement of a precise biological description (i.e. values for *α, β, λ* and *k* correspond to biophysical processes); b) incapacity to describe the diversity of tumor types (i.e. each is priming on a type of tumor), and c) the small amount and irregular sampling of the data. Usually, tumor growth data is small, only a few data points with, typically, days level granularity [18] and irregular spacing among measurements [19]. Moreover, the data has high variability due to: within tumor types specifics, chemotherapy effect on tumor growth [20], and heterogeneous measurement types (e.g. bio-markers, fMRI, fluorescence imaging [21], flow cytometry, or calipers [22]).

### C. Models of chemotherapy pharmacokinetics

Pharmacokinetics describe the distribution of chemotoxins in the body and their effects. From the point of view of a particular drug, the body can be thought of as comprising one or more compartments, each of which can be considered to be a space throughout which the substance is uniformly distributed, and has uniform kinetics of distribution or transport [23].

Moreover, pharmacokinetic modelling is a useful tool to describe and investigate the effect of covariates in drug variation. A number of population pharmacokinetic models have described the pharmacokinetics of the taxanes drug family [24]. More precisely, they addressed Paclitaxel monotherapy and have provided important insight into Paclitaxel pharmacokinetics - for more information see [25]. The importance of such a model was described by Norton and Simon which postulated that kinetic resistance of tumors can be counteracted through modulating dose density [10] but also dose adjustment in patients with hepatic impairment, suggesting, for example, that no dose adjustments are needed in patients with mild to moderate renal impairment [26].

In our study, we use the data from the computational model of intracellular pharmacokinetics of Paclitaxel of [17] due to its wide use in breast cancer chemotherapy schemes. The model describes the factors that determine the kinetics of Paclitaxel uptake, binding, and efflux from cells:

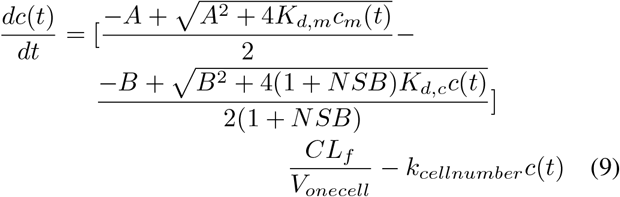

where:*V*_*onecell*_ is the average cell volume, *ICN* is the initial cell number at time 0, *NSB* is the proportionality constant for nonsaturable binding sites in cells, *k*_*cellnumber*_ is the rate constant for changes in cell number (i.e. depends on the pharmacological effect of Paclitaxel), *A* is a function of the constant for drug binding to proteins in medium *K*_*d,m*_ and *B* is a function of the constant for drug binding to proteins in cells, *CL*_*f*_ is the clearance of free drug by passive diffusion, on a per cell basis, and *c*_*m*_ concentration of drug in the medium, calculated as:

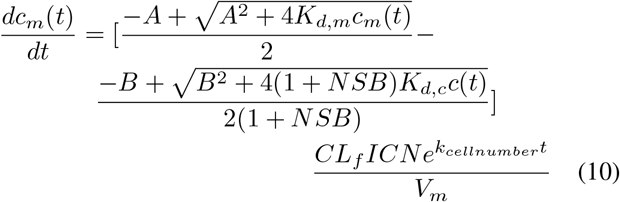

The intracellular pharmacokinetic model was used in [17] and in our study to predict the effect of cell density on drug accumulation with the mean value of experimental parameters chosen as: *K*_*d,m*_ = 781*nM, K*_*d,c*_ = 4.93*nM, NSB* = 0.148, *CL*_*f*_ = 3.3410^− 3^*ul/h/cell, k*_*cellnumber*_ = 0.0046*h*^− 1^ (see [17], Table 3). The model predictions were then compared with experimental results to evaluate the validity of the model.

Changes in cell number were represented by changes in volume which: 1) increased with time at low initial total extracellular drug concentrations due to continued cell proliferation and 2) decreased with time at high initial total extracellular drug concentrations due to the antiproliferative and/or cytotoxic drug effects, as reported in [17].

Such nonlinear effects are patient specific and parametrizing the model needs very detailed biological specification and analysis, which in vivo might not be feasible. Another challenge regarding the clinical use of Paclitaxel is the identification of optimal treatment drug administering schedules. The difficulty is in part due to the lack of a precise understanding of individual pharmacokinetics of Paclitaxel, i.e., drug effect as a function of drug concentration and treatment duration for each patient. Such challenges motivated our study.

### D. Objectives of the study

The objectives of our study are to demonstrate how CHIMERA, a combination of machine learning and mechanistic modelling can predict the chemotherapy and surgery sequencing, through:

- learning the tumor growth model *f* (*V*) from tumor growth data of breast cancer, and
- learning the pharmacokinetics *P* (*t, V*) of the chemotoxic dose response in the sequencing scheme,

for a truly personalized intervention in breast cancer patients.

## II. Materials and methods

In the next section we introduce the underlying mechanisms of CHIMERA as well as the experimental procedures used in our experiments.

### A. Introducing CHIMERA

CHIMERA is an unsupervised machine learning system based on Self-Organizing Maps (SOM) [27] and Hebbian Learning (HL) [28] used for extracting underlying functional relations among correlated timeseries describing therapy variables, such as tumor growth and chemotherapy pharmacokinetics. We introduce the basic mechanisms in CHIMERA through a simple example in Figure 1. Here, we consider data from a breast cancer growth function under sequential chemotherapy carried over 150 weeks from [29]. The two input timeseries (i.e. the number of tumor cells and the irregular measurement index over the weeks) follow a cubic dependency, depicted in Figure 1b-left.

**Fig. 1.**
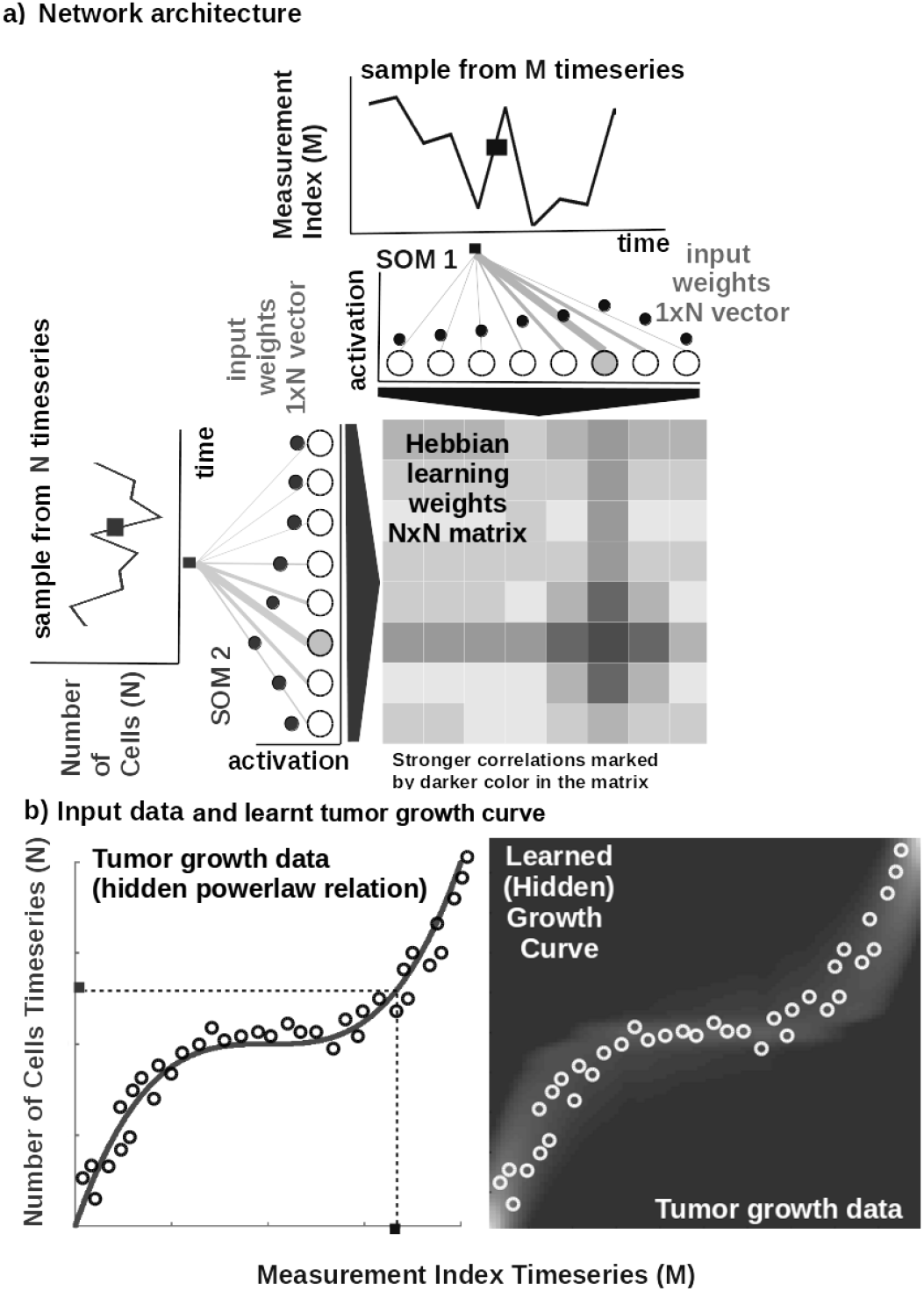
Basic functionality of CHIMERA: a) Basic architecture of CHIMERA: 1D SOM networks with *N* neurons encoding the timeseries (i.e. number of cells vs. measurement index), and a *NxN* Hebbian connection matrix coupling the two 1D SOMs that will eventually encode the relation between the timeseries, i.e. growth curve. b) Left: Tumor growth data resembling a non-linear relation hidden in the timeseries (i.e. number of cells vs. measurement index) Data from [29]. Right: Learnt tumor growth curve.

#### Core model

The input SOMs (i.e. 1D lattice networks with *N* neurons) encode timeseries samples in a distributed activity pattern, as shown in Figure 1a. This activity pattern is generated such that the closest preferred value of a neuron to the input sample will be strongly activated and will decay, proportional with distance, for neighbouring units. The SOM specialises to represent a certain (preferred) value in the timeseries and learns its sensitivity, by updating its tuning curves shape. Given an input sample *s*^*p*^(*k*) from one timeseries at time step *k*, the network computes for each *i*-th neuron in the *p*-th input SOM (with preferred value 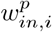 and tuning curve size 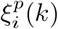) the elicited neural activation as

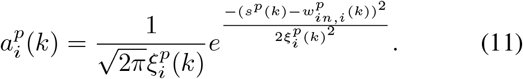

The winning neuron of the *p*-th population, *b*^*p*^(*k*), is the one which elicits the highest activation given the timeseries sample at time *k*

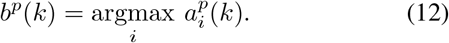

The competition for highest activation in the SOM is followed by cooperation in representing the input space. Hence, given the winning neuron, *b*^*p*^(*k*), the cooperation kernel,

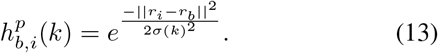

allows neighbouring neurons (i.e. found at position *r*_*i*_ in the network) to precisely represent the input sample given their location in the neighbourhood *σ*(*k*) of the winning neuron. The neighbourhood width *σ*(*k*) decays in time, to avoid twisting effects in the SOM. The cooperation kernel in Equation 13, ensures that specific neurons in the network specialise on different areas in the input space, such that the input weights (i.e. preferred values) of the neurons are pulled closer to the input sample,

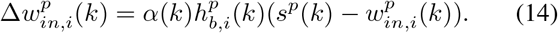

Neurons in the two SOMs are then linked by a fully (all-to-all) connected matrix of synaptic connections, where the weights in the matrix are computed using Hebbian learning. The connections between uncorrelated (or weakly correlated) neurons in each population (i.e. *w*_*cross*_) are suppressed (i.e. darker color) while correlated neurons connections are enhanced (i.e. brighter color), as depicted in Figure 1b-right. Formally, the connection weight 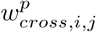 between neurons *i, j* in the different input SOMs are updated with a Hebbian learning rule as follows:

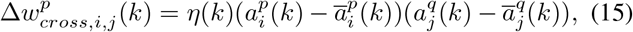

where 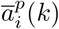 is a “momentum” like exponential moving average. Hebbian learning ensures that when neurons fire synchronously their connection strengths increase, whereas if their firing patterns are anti-correlated the weights decrease. The weight matrix encodes the co-activation patterns between the input layers (i.e. SOMs), as shown in Figure 1a, and, eventually, the learned growth law (i.e. functional relation) given the timeseries, as shown in Figure 1b-right.

Self-organisation and Hebbian correlation learning processes evolve simultaneously, such that both the representation and the extracted relation are continuously refined, as new samples are presented. This can be observed in the encoding and decoding functions where the input activations are projected though *w*_*in*_ (Equation 11) to the Hebbian matrix and then decoded through *w*_*cross*_.

#### Parametrization and read-out

In all of our experiments the data is fed to the CHIMERA which encodes each timeseries in the SOMs and learns the underlying relation in the Hebbian matrix. The SOM neural networks are responsible of bringing the timeseries in the same latent representation space where they can interact (i.e. through their internal correlation). In our experiments, each of the SOM has *N* = 100 neurons, the Hebbian connection matrix has size *NxN* and parametrization is done as: *alpha* = [0.01, 0.1] decaying, 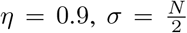 decaying following an inverse time law. We use as decoding mechanism an optimisation method that recovers the real-world value given the self-calculated bounds of the input time-series. The bounds are obtained as minimum and maximum of a cost function of the distance between the current preferred value of the winning neuron (i.e. the value in the input which is closest to the weight vector of the neuron in Euclidian distance) and the input sample at the SOM level.

### B. Datasets

In our experiments we used publicly available tumor growth datasets (see Table II), with real clinical tumor volume measurements, for different cell lines of breast cancer. This choice is to probe and demonstrate the versatility of CHIMERA in learning from tumor growth patterns induced by different types of cancer. For the pharmacokinetics of the chemotoxic drug (i.e. Paclitaxel), we used the data from [17] describing intracellular and extracellular concentrations of Paclitaxel during uptake. MCF7 breast cancer cells were incubated with 1 to 1000 nM Paclitaxel. The concentration of Paclitaxel in cells and culture medium were monitored for 24 h (see Table 1 and Table 2 in [17]. The volume of MCF7 cells in the exponential growth phase was determined using microscopic imaging (i.e. resolution of 164 × 200 px), where maximum (L) and minimum (W) diameters of a cell, were determined by counting the number of pixels and converting the value to micrometers 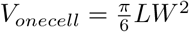.

**TABLE II.** Description of the datasets used in the experiments.

| Experimental dataset setup |  |  |  |  |  |
| --- | --- | --- | --- | --- | --- |
| Dataset | Cancer Type | Data Type | Data Points | Data Freq. | Source |
| 1 | Breast (MDA-MB-231 cell line) | Fluorescence imaging | 7 | 2x/week | [21] |
| 2 | Breast (MDA-MB-435 cell line) | Digital Caliper | 14 | 2x/week | [19] |
| 3 | Breast (MCF7, T47D cell lines) | Caliper | 8 | 1x/week | [30] |
| 4 | Breast (LM2-4LUC+ cell line) | Digital Caliper | 10 | 3x/week | [31] |

### C. Procedures

In order to reproduce the experiments in our study, the MATLAB ^®^ code and copies of all the datasets are available on GITLAB ^1^. Each of the three mechanistic tumor growth models (i.e. Logistic, Bertalanffy, Gompertz) and CHIMERA were presented the tumor growth data in each of the four datasets. When a dataset contained multiple trials and patients, a random one was chosen for testing and evaluation.

#### Mechanistic models setup

Each of the three ODE tumor growth models and the drug pharmacokinetic model was implemented as ordinary differential equation (ODE) and integrated over the dataset length. We used a solver based on a modified Rosenbrock formula of order 2 that evaluates the Jacobian during each step of the integration. To provide initial values and the best parameters (i.e. *α, β, λ, k*) for each of the four models the Nelder-Mead simplex direct search (i.e. derivative-free minimum of unconstrained multi-variable functions) was used, with a termination tolerance of 10*e*^− 4^ and upper bounded to 1000 iterations. Finally, fitting was performed by minimizing the sum of squared residuals (SSR).

#### CHIMERA setup

For CHIMERA the data was normalized before training and de-normalized for the evaluation. The system was comprised of two input SOMs, each with *N* = 50 neurons, encoding the volume data and the irregular sampling time sequence, respectively. Both input density learning and cross-modal learning cycles were bound to 100 epochs. The full parametrization details of CHIMERA are given in Parametrization and read-out section.

## III. Results

In the current section, we present the experimental results of our study and demonstrate that CHIMERA is capable to: a) learn the tumor growth model *f* (*V*) from tumor growth data of breast cancer, b) learn the pharmacokinetics *P* (*t, V*) of chemotoxic drug dose and c) to use the learnt quantities to provide a data-driven sequencing scheme for chemotherapy and surgery in a personalized breast cancer intervention.

### A. Learning the tumor growth function f (V)

The first experiment addressed the capability to learn the tumor growth model *f* (*V*) from data without imposing biological constraints upon the cell line, tumor size, number of cells etc. We demonstrate the superior learning capabilities of CHIMERA on the four publicly-available brest cancer clinical datasets described in Table II. As we can observe in Figure 2, CHIMERA learns a superior fit (i.e. lowest Sum Squared Error (SSE), Root Mean Squared Error (RMSE) and symmetric Mean Absolute Percentage Error (sMAPE)) to the tumor growth data, despite the limited number of samples (i.e. 7 data points for MDA-MD-231 cell line dataset and up to 14 data points for MDA-MD-435 cell line dataset). Despite their ubiquitous use, the classical tumor growth models (e.g. Gompertz, von Bertalanffy, Logistic) are confined due to: a) the requirement of a precise biological description - as one can see in the different sigmoid shapes in Figure 2; b) incapacity to describe the diversity of tumor types - as shown in the across-cell line summary statistics in Figure 3 and; c) the small amount and irregular sampling of the data - visible in the relatively poor fit to the data, captured in Figure 2.

**Fig. 2.**
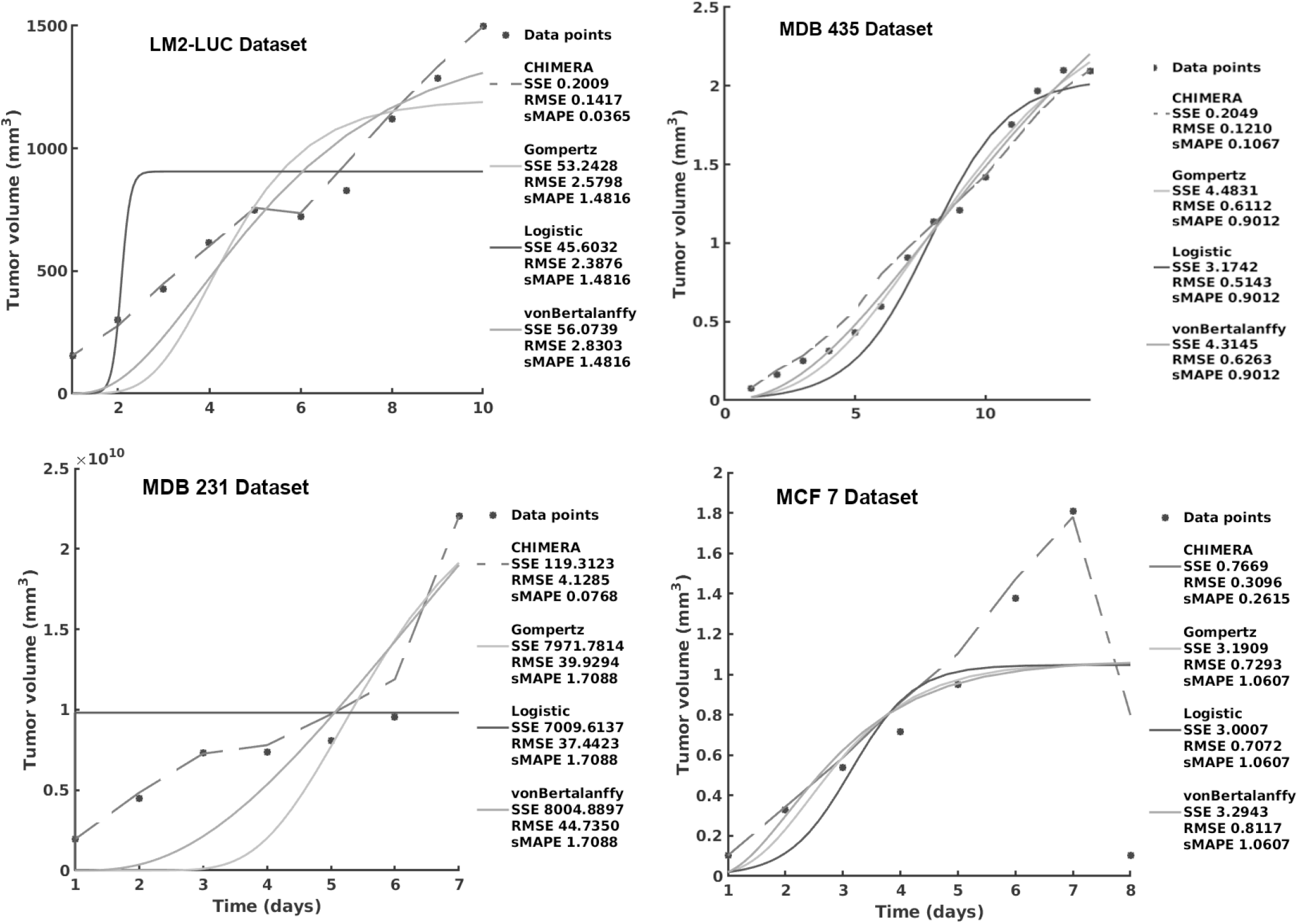
Evaluation of the tumor growth models on the different datasets: accuracy evaluation. CHIMERA is decoded from the learnt Hebbian weight matrix. The decrease in the MCF7 Dataset is due to the administered chemotherapy and demonstrates the adaptivity of CHIMERA in capturing growth behaviours.

**Fig. 3.**
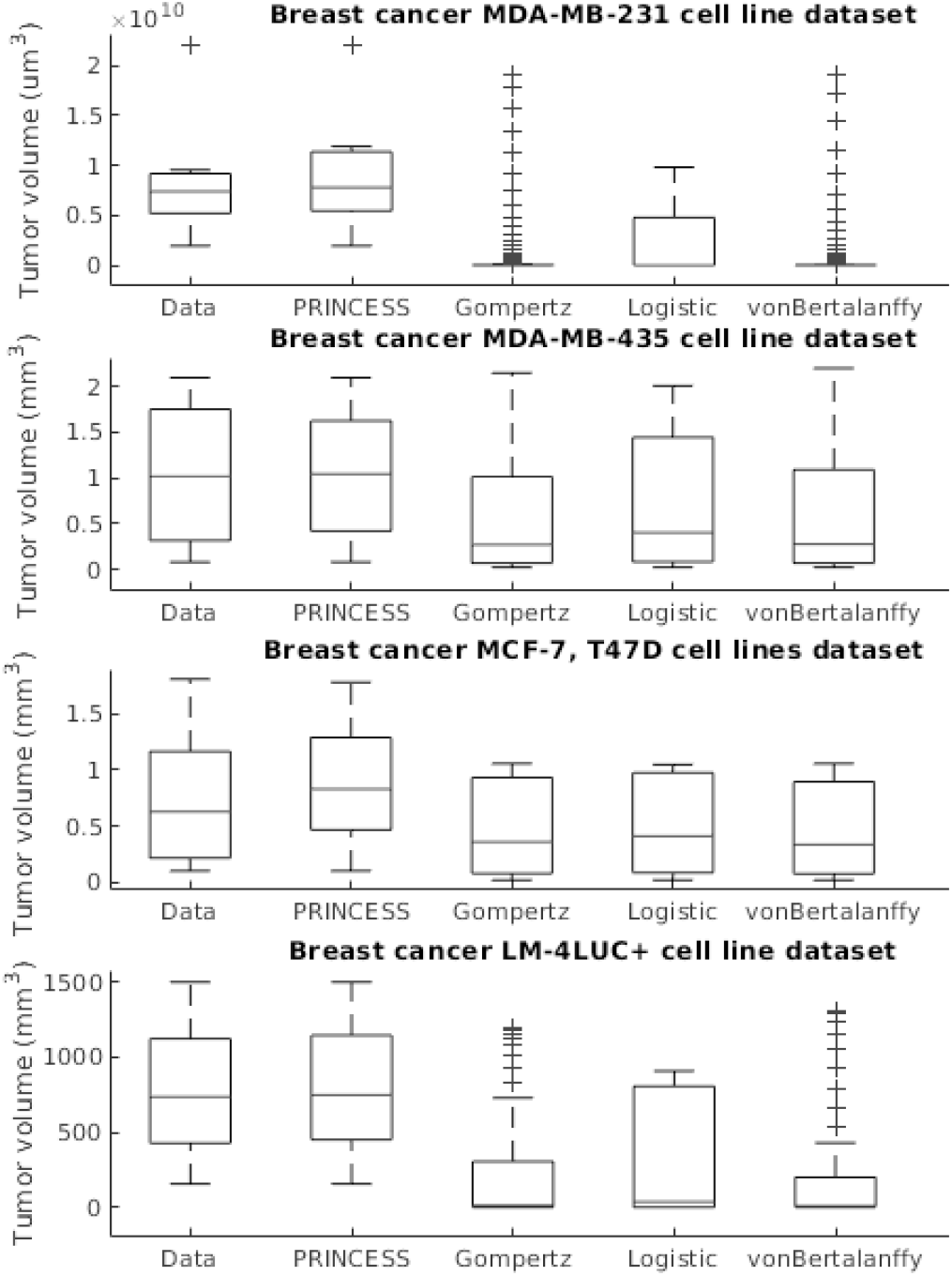
Evaluation of the tumor growth models on the different datasets: summary statistics.

### B. Learning the pharmacokinetics P(t, V)

The second experiment focused on extracting the pharmacokinetics of a chemotoxic drug typically used in breast cancer, namely Paclitaxel. Its pharmacokinetics function *P* (*t, V*) is dependent of its concentration concentration *c*(*t*) at time *t* and *V* the volume of the tumor. In our experiments, the drug concentration *c*(*t*) has two components as following the model in [17], namely the intracellular and extracellular Paclitaxel (Equations 9 and 10). Due to the multiple empirical parameters the model needs and the relatively small dataset, parametrizing the model in Equations 9 and 10 requires many biological assumptions that typically do not hold in vivo. Such variability is typically not captured by such models, hence failing to actually describe the pharmacokinetic behavior in a certain patient.

### 1) Cellular concentration

As one can see in Figure 4, the intracellular concentration kinetics of Paclitaxel is highly nonlinear. CHIMERA is able to extract the underlying function describing the data without any assumption about the data and other prior information, opposite to the model from [17] which used Equation 7 to fit the data. Interestingly, CHIMERA captured a relevant effect consistent with multiple Paclitaxel studies [25]. Namely, that the intracellular concentration increased with time and approached plateau levels, with the longest time to reach plateau levels at the lowest extracellular concentration - as shown in Figure 4.

**Fig. 4.**
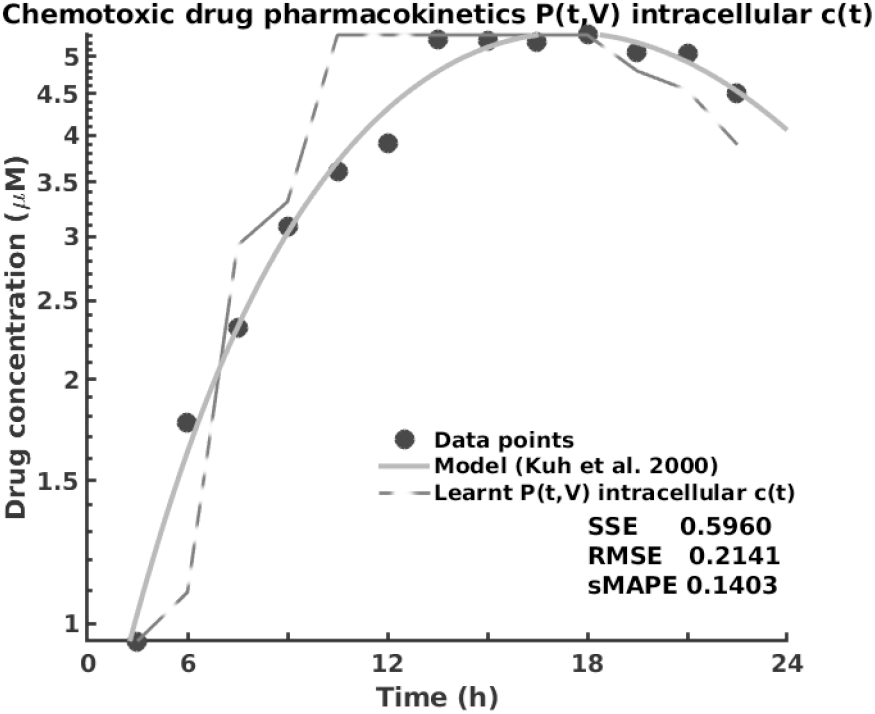
Learning the pharmacokinetics *P* (*t, V*) of the intracellular Paclitaxel concentration *c*(*t*). Data from [17] and model in Equation 7, log scale plot.

### 2) Extracellular concentration

Analysing the extracellular concentration in Figure 5, we can see that CHIMERA extracted the trend and the individual variation of drug concentration after the administration of the drug (i.e. in the first 6h) and learnt an accurate fit without any priors or other biological assumptions. Interestingly, CHIMERA captured the fact that the intracellular drug concentration increased linearly with extracellular concentration decrease, as shown in Figure 4 and Figure 5.

**Fig. 5.**
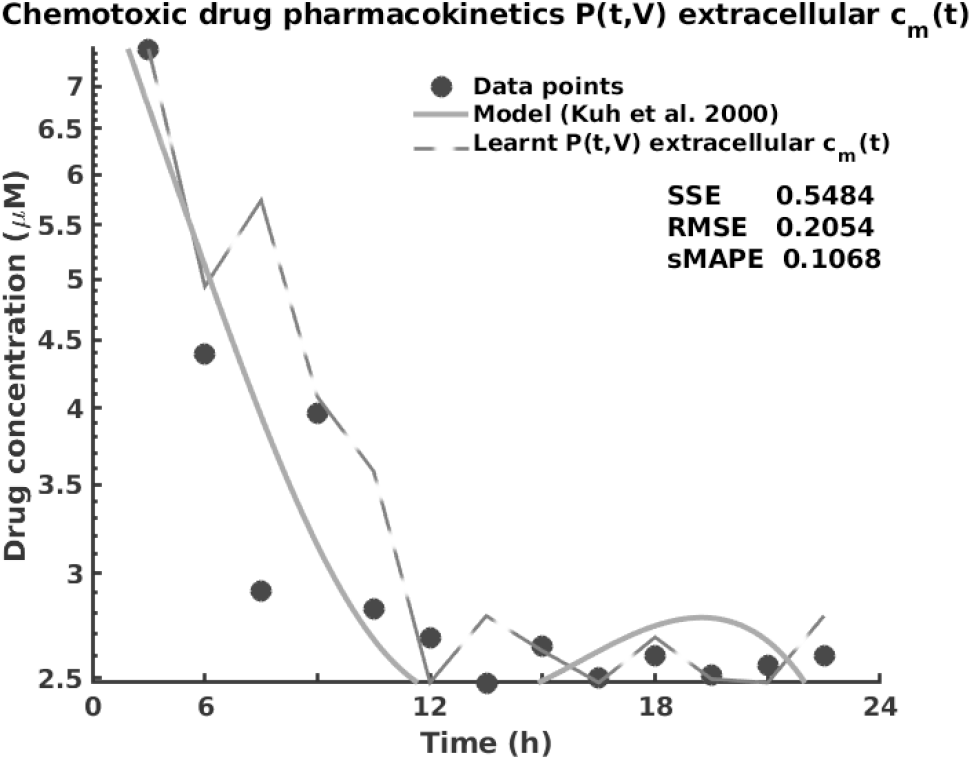
Learning the pharmacokinetics *P* (*t, V*) of the extracellular Paclitaxel concentration *c*_*m*_(*t*). Data from [17] and model in Equation 8, log scale plot.

### C. Chemotherapy-Surgery Sequencing

In this sub-section we combine all the results in the previous sub-sections on learning the tumor growth function *f* (*V*) and the pharmacokinetics *P* (*t, V*) with a mechanistic modelling framework that demonstrates the capabilities of CHIMERA in sequencing chemotherapy and surgery in breast cancer. We evaluate the potential sequencing under the two hypotheses, log-kill and Norton-Simon, respectively. In order to simplify the formulation, we consider the tumor volume *V* = *Nυ*, where *N* is the number of cells and *υ* is a constant describing cell volume and the volume of intercellular space. We assume, for simplicity, that the growth model follows a Gompertz growth curve with a transport constant 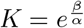,

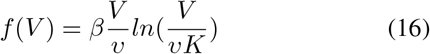

and the pharmacokinetics function is

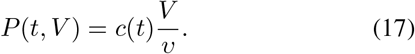

Following our derivation in Section, 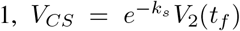 and *V*_*SC*_ = *V*_1_(*t*_*f*_) corresponding to tumor sizes in neo-adjuvant and adjuvant sequences, respectively. Under the log-kill assumption if we let 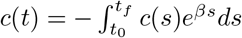 then

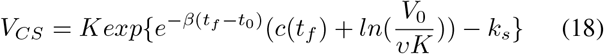

and

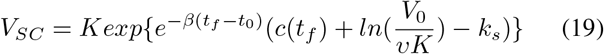

where 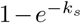, with *k*_*s*_ a constant, is the fraction of the tumor that is removed. Therefore,

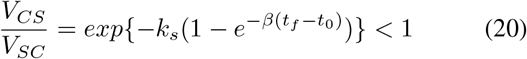

hence *V*_*CS*_ *< V*_*SC*_. Under the Norton-Simon assumption, if we let 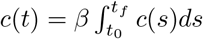, then

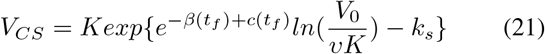

and

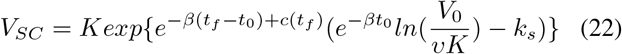

where 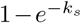, with *k*_*s*_ a constant, is the fraction of the tumor that is removed. Therefore,

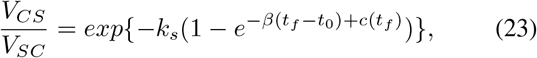

which for 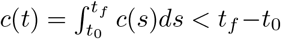 determines *V*_*CS*_ *< V*_*SC*_. Similarly, one can derive the sequencing for the Logistic and von Bertalanffy models by replacing Equation 16 with the corresponding equations in TableI-B. In order to evaluate the sequencing capabilities of CHIMERA with respect to traditional biologically parametrized models (we only present Gompertz in Table III, analog results for Logistic, von Bertalanffy), we consider the dataset of breast cancer (MCF7 cell line) from [30] described in our Experimental setup. We use the derivations in Equations 6 and 3 and fill in with the decoded values from the learnt tumor growth (Figure 2) and learnt pharmacokinetics (Figures 4 and 5). Without paying the price of extensive parametrization and biological dependency, CHIMERA uses learnt tumor growth and pharmacokinetics to infer the most appropriate sequence of therapy, consistent with its mechanistic counterparts, as shown in Table III.

**TABLE III.** Evaluation of the chemotherapy-surgery sequencing prediction.

| Model | Log-kill hypothesis | Norton-Simon hypothesis | Biological Parameters |
| --- | --- | --- | --- |
| Gompertz | $V_{CS} < V_{SC}$ | $V_{CS} < V_{SC}$ if $c(t) = \int_{t_0}^{t_f} c(s)ds < t_f - t_0$ | $\beta, K, v$ and Eq.s 7, 8 |
| CHIMERA | $V_{CS} < V_{SC}$ | $V_{CS} > V_{SC}$ | none |

## IV. Conclusion

As ever, there will be no “one-size-fits-all” treatment for breast cancer and the focus should be on optimising patient characterization. This can be achieved through a data-driven approach in which individual patient data describing tumor growth (e.g. histology, imaging) and chemotoxic drug effect (i.e. pharmacokinetics, drug interactions) are used in combination to extract the optimal sequence of therapy. CHIMERA is an initial effort to offer a personalized solution in chemotherapy-surgery sequencing that can handle biological variability of tumors, the limited size of patient data, and the variability in chemotoxic drug response, in a data-driven way. We demonstrated and evaluated CHIMERA’s capabilities in a series of experiments that emphasize the need for combining data-driven and mechanistic modelling in oncology.

## Footnotes

1 https://gitlab.com/akii-microlab/ismco2020

